# In vivo timelapse imaging and analysis of Golgi satellite organelle distribution and movement in the neural progenitor cells of the brain

**DOI:** 10.1101/2024.02.16.580740

**Authors:** Lindsay D Arellano, Jennifer E Bestman

## Abstract

The dividing stem cells of the developing brain are the radial glial neural progenitor cells (NPCs), multifunctional cells that proliferate to generate all of the cells of the brain, but also act as scaffolds for their migrating neuron progeny, guideposts for pathfinding growing axons and regulators of synaptic activity. These remarkable cells perform these very different activities while remaining in contact with the inner and outer surface of the ever-growing brain. NPCs synthesize proteins locally to support the compartmentalized protein expression required for the cells to perform their specialized functions, but it is not clear how the necessary processing that normally occurs in the Golgi apparatus is achieved at locations far from the cell body. Golgi satellites, motile organelles and members of the protein maturation machinery, control protein glycosylation and maturation in polarized cells like neurons. To investigate whether NPCs also rely on Golgi satellites, we expressed a fluorescent reporter to label Golgi satellites in the NPCs in the intact brains of *Xenopus laevis* tadpoles. Quantitative analysis of *in vivo* timelapse images revealed dynamic, motile Golgi satellites that distribute throughout the cell, suggesting that NPCs have local proteostasis to support their diverse functions.

## 1. Introduction

Radial glial neural progenitor cells (NPCs) are dividing stem cells in the developing vertebrate brain. Also known as radial astrocytes, they have a periventricular-positioned cell body at the center of the brain, and a radial process (aka, basal process) that projects from the cell body and terminates in a flattened basal endfoot that interfaces with the blood brain barrier (1, 2). The tissue-spanning morphology of NPCs grows with brain development, reaching thousands of microns in some animals (3). Whether NPCs divide symmetrically to expand the pool of progenitors or divide asymmetrically to generate a neuron with each cell division, typically just one of the cellular progeny retains the radial process and the other newborn cell must elaborate its own radial process or begin the development of its dendritic arbour and axon (4). These complicated cell divisions occur at a rate of +1000/second in the development of the human brain (5). Given the number of developmental disorders that occur when cell proliferation goes awry (6), there is a great need to understand the fundamental cell biology of these important cells.

To regulate proteins efficiently and responsively, polarized cells in a variety of organisms rely on local protein synthesis whereby mRNAs, ribosomes and “satellite” protein maturation machinery transported to distant sites so that proteostasis can be controlled independently from the main machinery in the cell body (7-9). Local protein synthesis is particularly critical for neurons that have complex dendritic branches and thousands of synaptic contacts that must be independently regulated (9) and it has similar roles in the complex architecture of astrocytes (10). NPCs also exhibit regional differences in gene expression and function. The apical and basal compartments of NPCs have different transcript expression (11), different levels of local protein synthesis (12), resulting in distinct subcellular protein localization (13, 14). NPCs display spatially restricted calcium transients (15, 16) and filopodial branch movements (17). The specialized morphology of NPCs provides a scaffold for migrating newborn neurons in the developing cortex (18), and their basal endfeet are essential guides for retinal ganglion cell axons as they enter the brain (19). At the end of embryonic development NPCs lose their radial morphology and terminally differentiate into multipolar astrocytes where local translation is critical how these macroglia regulate synapses and maintain the blood brain barrier in the adult brain (20).

We were interested in describing whether NPCs have Golgi satellite networks distributed away from the main protein sorting Golgi complex machinery housed in their somata to support the functional and spatial specializations of these cells. In neurons, Golgi microsecretory organelles inhabit distal branches where they function as acentrosomal microtubule organizing centers that regulate dendritic cytoarchitecture (21) and participate in cargo trafficking and protein glycosylation (9). The fragmentation of Golgi and formation of Golgi satellites that disperse from the cell body into the dendritic arbour is linked to neural activity (22). Neurons utilize ER export sites and Golgi satellites to traffic supplies for formation of plasma membranes, facilitating asymmetrical growth in dendrites (23-25), and Golgi satellites alter the glycoproteome of the dendritic membrane (26, 27). Proper control of the glycoproteome has implications in molecular interactions, cell-cell adhesion, and neuronal differentiation and function (28). Without the need to rely on one centralized perinuclear Golgi apparatus, these satellite organelles allow neurons to independently control proteostasis at distant positions. The main Golgi apparatus has an unusual morphology and localization in the NPCs (29), suggesting that NPCs would also employ alternatives to typical Golgi processing. Despite the recent interest in Golgi satellites and local control of proteostasis, it is currently not clear if the progenitors to neurons, the NPCs, also have Golgi satellites.

Here we use *in vivo* 4D microscopy to investigate Golgi satellites in the NPCs in the brain of *Xenopus laevis* tadpoles where transfection and time lapse imaging can be conducted in the intact animal. To label the NPCs and their Golgi satellites, we modified pGolt3, a fluorescent reporter that efficiently and specifically labels the Golgi satellites (25, 30). We expressed pGolt3 in *Xenopus* NPCs revealing that, like neurons, the trans-Golgi-network is decentralized and dispersed in NPCs. Our morphometric analyses of 3D timelapse images reveal that NPCs contain dynamic, motile Golgi satellite organelles that evenly distribute throughout the cells. Our results suggest that NPCs, the essential stem cells of the developing brain, may rely on a distributed Golgi microsecretory network to support proteostasis.

## 2. Materials and Methods

### (a) Plasmid Construction

The pCMV-eGFP-p2A-Golt3 expression vector was assembled by inserting cytosolic eGFP fused to the p2a peptide (31); a gift from Philip Washbourne (Addgene plasmid # 80807) into the pGolt3 plasmid (a gift from Michael Kreutz, Addgene plasmid # 73297). The original pGolt3 vector encodes the mCherry red fluorescent protein fused to endoplasmic reticulum export signal sequences from Scap, a Golgi export protein, and the transmembrane domain of the calneuron-2, a protein enriched in the Golgi (30). We assembled this plasmid by linearizing pGolt3 with NheI and AgeI restriction enzymes and inserting eGFP-P2a fragment (Forward primer: agcgagtcagtgagcgag Reverse primer: gctACCGGTaagaaagctgggtaaggac) generated from the pME-GFP-p2a template. Our insertion of the “self-cleaving” P2A peptide between eGFP and mCh-Golt3 produces independent two proteins under the control of the cytomegalovirus (CMV) promoter (31); (figure 1).

**Figure 1.**
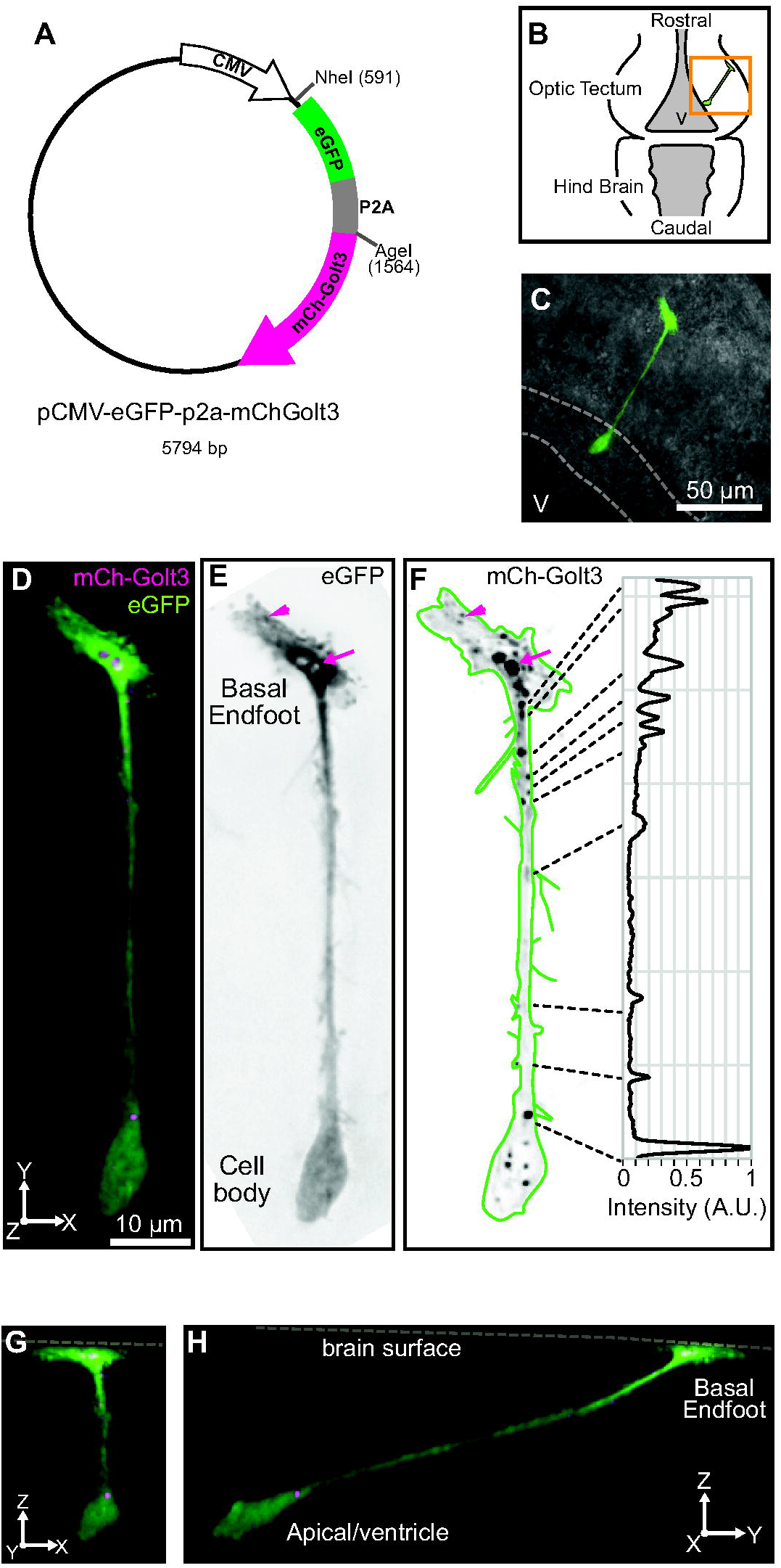
Golgi satellite plasmid vector and its expression in NPCs. A) Map of plasmid reporter used to express cytosolic eGFP and the Golgi satellite marker, mCherry-Golt3. B) Diagram of the NPCs and the *Xenopus* brain. C) a labelled NPC in the tectum (v: ventricle). D-F) z-projection of the fluorescently-labelled NPC with each channel displayed as inverted greyscales. Inset (f) is the plot profile of the mCh-Golt3 fluorescence with the locations of Golgi satellites. G and H) are y- and x-oriented projections of the NPC revealing its large basal endfoot and radial morphology.

### (b) Tadpole rearing and In Vivo Electroporation

Albino *Xenopus laevis* tadpoles were purchased from *Xenopus express* (Xenopus.com) as embryos and reared in HEPES-buffered Steinberg’s solution (58.0 mM NaCl, 0.67 mM KCl, 0.34 mM Ca(NO_3_)_2_, 0.83 mM MgSO_4_, 3.0 mM HEPES, pH 7.2) at 23°C under a 12h light/12h dark cycle. Tadpoles were fed *Xenopus express* Premium Tadpole Food beginning at stage 46. After the first confocal image, the tadpoles were housed singly in 6-well cell culture plates. The animal protocol was approved by the Institutional Animal Care and Use Committee of William & Mary in accordance with NIH guidelines.

NPCs of stage 46 (32) tadpoles were transfected using *in vivo* electroporation. After anesthetizing the tadpoles with 0.02% MS-222 in Steinburg’s solution, they were positioned under a dissecting stereomicroscope. The pCMV-eGFP-p2a-Golt3 plasmid (500 ng/µL) mixed with 1% Fast Green dye was loaded into a pulled microcapillary glass pipette. Using a micromanipulator, millisecond pressure pulses injected the plasmid solution into brain ventricle. Custom platinum electrodes were placed on the surface of the head lateral to the optic tectum using a micromanipulator, and 4 voltage pulses (1.6ms duration at 35V pulse transformed to an exponential waveform with a 3µF capacitor) were applied and then repeated with reversed polarity to electroporate the tectal NPCs. Afterward, the tadpoles were transferred to recover in Steinberg’s solution and images were acquired 24-36 hours later.

### (c) Time Lapse Fluorescence Confocal Microscopy and Image Analysis

Tadpoles were anesthetized in 0.02% MS-222 in Steinburg’s media and placed in a custom Sylgard form so the dorsal surface of their head can be topped directly with a glass coverslip and positioned under the microscope objective. Images of the labelled NPCs were acquired using a VIVO spinning disc confocal system (3i, Inc.) composed of a Zeiss Examiner microscope, a water immersion objective (Zeiss LD C-Apochromat 63x/1.15 W), a Yokogawa CSU-X spinning disc confocal and Photometrics Prime 95B camera. Full 3D confocal stacks (z-depth between 50 and 150µms) were acquired at 0.36µm z-step intervals of the eGFP + mCh-Golt3 signals before rapid 3D timelapse images (30 second intervals at 1µm z-steps) to minimize harm from the excitation light and to speed image acquisition followed by a final image of the eGFP and mCherry signals.

3D confocal stacks of the NPCs were cropped and projected to produce the 2D images, then converted to 8-bit RGB images using Slidebook 6.0 software (3i, Inc.). To quantify the distribution and movement of the mCh-Golt3 signal, the original 16 bit timelapse images were opened with FIJI/Image J (33), and the radial process was reconstructed in 3D using the Simple Neurite Tracer tool (SNT; (34). We generated straightened representations of the radial process and used Multikymograph to produce kymographs.

Movement of the mCh-Golt3 labelled Golgi satellites was calculated as a cumulative change in intensity for each position of the cell’s radial process. We measured the intensity for each position over time normalized to the maximum before calculating the change in fluorescence intensity between each timepoint. The same analysis was performed on the cytosolic eGFP signal from the first and last images to derive a baseline value of fluorescence intensity fluctuations. Events when the mCh-Golt3 intensity changed more than the baseline cytosolic eGFP signal were recorded. The cumulative mCh-Golt3 change is the sum of change events for each position. To quantify the stability of mCh-Golt3, the intensity at each position was summed and normalized to the position with the greatest summed intensity, revealing the sites with the most stable and intense mCh-Golt3 signal. To compare these morphometric values across multiple cells, we normalized the lengths of the radial processes and then calculated the cumulative probability of mCh-Golt3 change or stability for each cell. Values are reported as means ± standard error of the mean; the lower and upper 95% confidence intervals (Cis) are reported in brackets. JMP17 statistical software was used to assess the significance of the regressions, and nonparametric Wilcoxon tests were used to detect differences in the proportion of mCh-Golt3 changes.

## 3. Results

### (a) The Golgi satellite mCherry-Golt3 reporter and its expression in neural progenitor cells

The brain of *Xenopus laevis* tadpoles lies directly beneath the animal’s transparent skin, allowing direct *in vivo* transfection of plasmid reporters and time-lapse imaging of the fluorescently labelled cells in the intact animal using 3D timelapse confocal microscopy (35). In this study we were interested in observing and quantifying Golgi satellite organelles in the radial glial neural progenitor cells (NPCs) of the optic tectum (figure 1b), the area of the brain that receives the axon inputs from the eye, controls visual behaviours and continues to show cell proliferation and sensory-dependent plasticity throughout development (36-38). To distinguish NPCs from the other cell types in the tectum, we modified the pGolt3 fluorescent reporter (a gift of Michael Kreutz) to create pCMV-eGFP-p2A-mCh-Golt3, which drives expression of cytosolic eGFP independently from the mCh-Golt3 Golgi satellite tag (figure 1A and methods).

Figure 1c shows the distinct slender morphology of a labelled NPC within the optic tectum tissue. The cell’s soma is at the ventricular surface deep in the brain (dotted lines, figure 1c) and its radial process extends dorso-laterally to terminate at the outer surface of the brain. The z-:projection of this cell is shown in figure 1d and this image, together with the y- (figure 1g) and x-oriented (figure1h) projections, reveal its nearly perfectly straight morphology and 20µm diameter basal endfoot that expands across the surface of the brain.

The cytosolic eGFP channel (figure 1e) and the mCh-Golt3 signal (figure 1f) are displayed as inverted grayscale images to better reveal the fine filopodial process (reconstructed as the green outline in figure 1f) and the variation of mCh-tagged Golgi satellite intensities (figure 1F). The inset in figure 1F is a profile of the mCh-fluorescence along the radial process with dotted lines connecting the positions of the Golgi satellite puncta to the graph of their relative intensities. These data show a roughly 4-fold difference in Golgi satellite signal intensities in this cell. The arrow and arrowhead in Figs 1e and 1f point to examples of Golt3-labeled Golgi satellites in the pial endfoot. These data show that like in neurons, NPCs also contain Golgi satellite organelles that disperse from the cell body and distribute throughout the NPCs occupying positions in the distal pial endfoot that are more than 80µm from the cell body.

### (b) Rapid timelapse of mCh-Golt3 labelled Golgi satellite movement in neural progenitor cells

We next acquired rapid 3D confocal timelapse images of the NPCs and the mCh-Golt3 reporter to quantify the degree to which the Golgi satellites move over time in these cells. Figure 2a-c are z-projections of the cytosolic eGFP and mCh-Golt3 signals together (figure 2a) and as inverted grayscale images (Figs 2b and 2c). This NPC was repeatedly imaged at 30 second intervals for 14.5 minutes, and the Figs 2d-g show the projected images captured at 0, 4, 10, and 14.5 minutes. We made 3D reconstructions of the cell at all timepoints to create the kymograph (figure 2h) of the mCh-Golt3 expression in the radial process over time.

**Figure 2.**
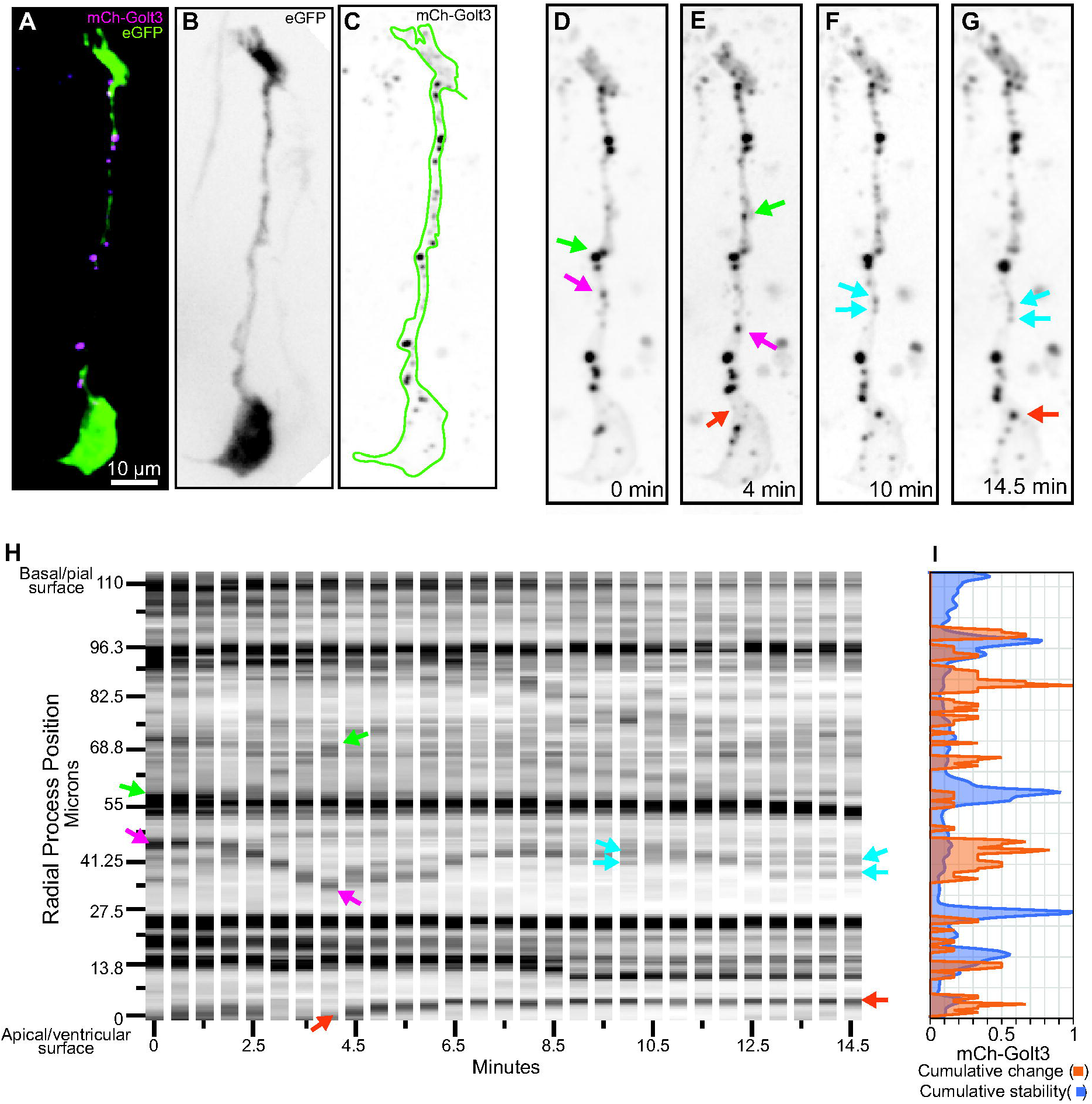
*In vivo* timelapse of mCh-Golt3 labelled Golgi satellites. A-C) Z-projection of NPC with greyscale images of the cytosolic eGFP (b) and mCh-golt3 expression (c). D-G) 4 timepoints of the timelapse. H) Kymograph of the mCh-Golt3 expression over time revealing movement of Golgi satellites. Arrows correspond between the images in D-G and the kymograph and point to moving Golgi satellites. I) Quantification mCh-Golt3 signal in this cell showing locations of change (orange) and locations of stationary expression (blue).

The arrows in Figs 2d-g correspond to arrows in the kymograph. The green arrows point to a mCh-Golt3-labeled Golgi satellite that, beginning at minute 2.5, moves 14µm away from the cell body until it can no longer be distinguished from the other tagged Golgi satellites in this area of the cell. Similarly, at minute 4, a Golgi satellite (orange arrow) leaves the soma and enters the radial process where it remains for the duration of the timelapse (orange arrow). Golgi satellites also show retrograde (soma-directed) movement as indicated with the magenta arrow which points to a Golgi satellite that moved ∼15µm between minute 2 and 4. This Golgi satellite then returns nearly to its starting position (cyan arrows) where at minute 10 it appears to disperse into multiple small puncta that can be distinguished at the final timepoint (cyan arrows).

The kymographs provide a sensitive visual representation of the mCh-Golt3 Golgi satellite movements for each cell, revealing that often the most dynamic Golgi satellites were small or expressed low levels of the mCh-Golt3 reporter. Because it is impossible to confidently identify and track many of these more dynamic Golgi satellites, we describe the relative stability and changes in mCh-Golt3 fluorescence that occur at each position along the radial process over the duration of the timelapse. From these data, we generated two morphometric measurements of mCh-Golt3 Golgi satellite dynamics: summed change per position and summed intensity per position (figure 2i). The summed change reflects the number of significant changes in mCh-Golt3 intensity at each position over the course of the timelapse; these data are presented relative to their maximum for this cell (figure 2i; orange). We find that the mCh-Golt3 signal was stable in 54% of the radial process, and single changes in mCh-Golt3 signal detected in 20% of the positions. We also totalled the intensity of the mCh-Golt3 signal for each position along the cell’s radial process and plotted these intensity values normalized to the largest value (figure 2i; blue). This summed intensity metric reflects the locations of the mCh-Golt3 labelled Golgi satellites that are stable throughout the timelapse session. These data provide a quantitative assessment of the mCh-Golt3 expression and show that stable sites and positions with more dynamic Golgi satellites are distributed evenly along the radial process of this NPC.

### (c) Morphometric quantification of mCh-Golt3 Golgi satellite distributions and movements

We next applied the summed change morphometric analysis to the five NPCs that completed the timelapse imaging protocol to compute average levels of motile mCh-Golt3-labeled Golgi satellites in these cells (figure 3a). We found no change in mCh-Golt3 signal in 32±8.0% [9.8-54.1%] of the NPCs’ radial processes, a proportion that is not significantly different from the number of positions that displayed a single change (32±6.8% [13.1-50.9%]) or the 22.4±5.0% [8.6-36.2%] of the positions along the radial processes that displayed 2 changes in the mCh-Golt3 signal over time. Positions with more mCh-Golt3 movement were rare and occurred significantly less than the single movement events (p=0.01), with only 7.6%±2.7% [3.6-17.1%] positions displaying 3 or more changes. These data indicate that the average mCh-Golt3-labeled Golgi satellite in NPCs are motile and will move from its initial position over the course of a 15-minute timelapse imaging session.

**Figure 3.**
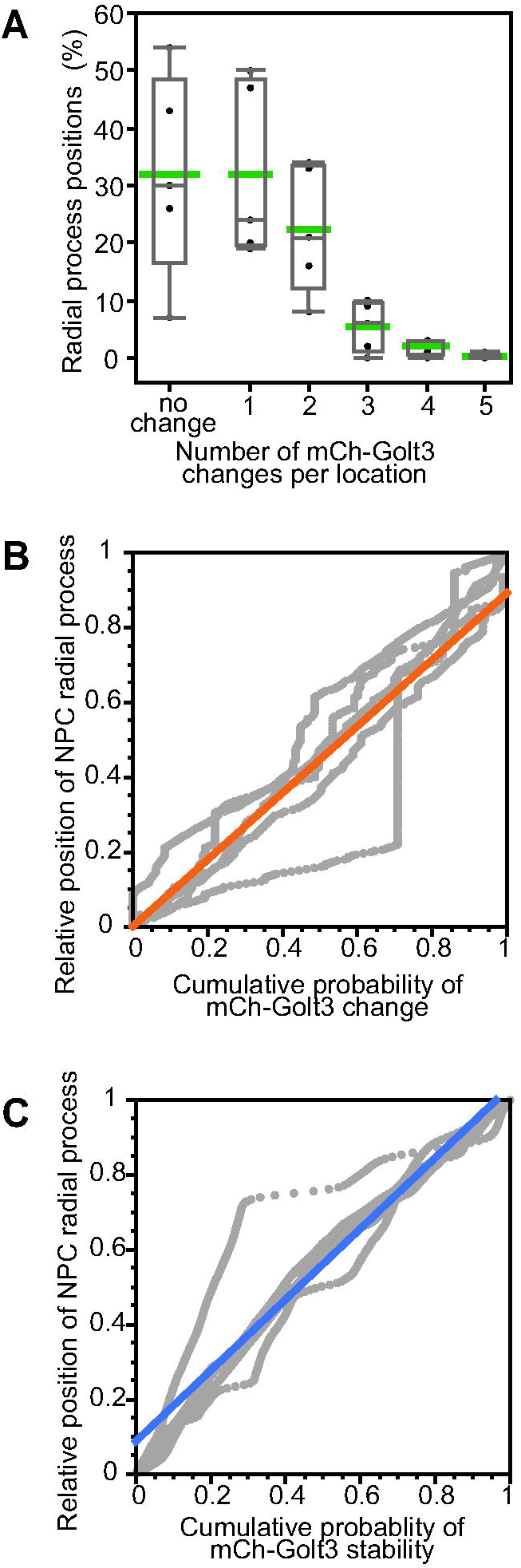
mCh-Golt3-labeled Golgi satellite timelapse quantification. A) Average number of changes in mCh-Golt3 expression that occurred across the NPC radial process over the entire timelapse session were calculated for 5 NPCs. Averages are shown in green along with the interquartile ranges and 95% CIs. B) Regression analyses show that the positions with changes in Golgi satellite expression (indicating movement) and locations of stable expression (C) are equally distributed along the apical-basal axis of NPCs.

To test whether there are regional or subcellular differences in the levels of mCh-Golt3 Golgi satellite movements in NPCs, we calculated the cumulative probability of intensity changes for all NPCs and plotted their distribution relative to their positions from the cell body (position = 0) to the pial endfoot (position = 1; figure 3b). The data for each NPC are shown in grey and the solid orange line is the linear fit for the data (R-square = 0.83). We found a significant positive correlation between the cell position and cumulative probability of mCh-Golt3 change (r=0.91 [0.90-0.92%], p<0.0001) indicating that movements of mCh-Golt3 were evenly distributed along the cells’ radial processes. In figure 3c, we similarly found a significant positive correlation between the cumulative probability of mCh-Golt3 summed intensities, a metric of stable Golgi satellites, and their positions in the NPCs (r= 0.96 [0.95-0.96%], p<0.001). The individual NPC values are shown in grey and are fit with the solid blue line (R-square = 0.91). Together the data in Figs 3b and c indicate that the mCh-Golt3-labeled Golgi satellites are evenly dispersed throughout the polarized morphology of the NPCs and that the movement of Golgi satellites is not associated with their proximity to the cell body or distal pial endfoot of the cells.

## 4. Discussion

Here we present the first description of the abundant and dynamic Golgi satellite microsecretory organelles in radial glial NPCs in the intact brain. Our analyses reveal that they are motile, moving in both anterograde and retrograde directions, but we find no differences in the subcellular distributions of the Golgi satellites in NPCs or regional differences in their likelihood to move. These data are similar to what has been shown in neurons (25) where the mCh-Golt3 tagged Golgi satellites distribute into even the most distal dendritic branches, and timelapse imaging and fluorescence recovery after photobleaching analysis show highly dynamic movement of the Golgi satellites and the organelles direct regulation of protein glycosylation far from the cell body.

There are clear connections between protein glycosylation and the healthy development and function of the brain: when the glycosylation of surface proteins in neurons and glia is misregulated, protein activity is altered and these changes are linked to many neurological disorders (39, 40). How protein glycosylation is regulated to support the precise cellular activity of neurons and glia is less clear. Recent data shows that the glycoprotein composition of neurons changes in response to neural activity, and that proteins trafficked through Golgi satellites have altered glycosylation (26). Our data suggests that the abundance and motility of Golgi satellites in progenitor cells of the brain may support protein sorting and cellular modifications, significantly contributing to the fate and function of NPCs.

## Acknowledgements

We gratefully acknowledge the support provided by the NICHD (R15HD099023) and the contributions of the 2021 Biol 406 class.

